# Aging-associated decline of phosphatidylcholine synthesis is a malleable trigger of natural mitochondrial aging

**DOI:** 10.1101/2024.04.25.591184

**Authors:** Tetiana Poliezhaieva, Pol Alonso Pernas, Lilia Espada, Melike Bayar, Yuting Li, Handan Melike Dönertaş, Maria A. Ermolaeva

## Abstract

Mitochondrial dysfunction is a prominent hallmark of aging contributing to the functional decline of metabolic plasticity in late life. While genetic distortions of mitochondrial integrity elicit premature aging, the mechanisms leading to “natural” aging of mitochondria are less clear. Here we initially used proteomics, genetics and functional tests in wild type *C. elegans* and long-lived *clk-1(qm30)* and *isp-1(qm150)* mitochondrial mutants to identify molecular pathways that safeguard longevity amid persistent mitochondrial inefficiency. Strikingly, these analyses and subsequent transcriptomic and functional tests in the human system revealed aging-associated decline of phosphatidylcholine (PC) synthesis as a trigger of mitochondrial network disruption, which contributes to mitochondrial dysfunction during normal aging. Moreover, we found that ectopic boosting of PC levels via diet restores late life mitochondrial integrity *in vivo* and rescues metabolic plasticity in cell culture tests. Our work thus uncovered a novel natural driver of mitochondrial aging that is malleable by dietary interventions.

## Introduction

Aging is characterized by the progressive deterioration of multiple cellular and organismal functions^1^, and one of key challenges in countering aging-associated functional decline is to identify and mitigate core mechanistic drivers of the aging process. Mitochondrial dysfunction is clearly one of the best recognized hallmarks of aging^1, 2^, and in addition to causing cellular deterioration in late life, it blunts the efficacy of anti-aging interventions that rely on metabolic plasticity such as dietary restriction (DR) and DR mimetics^3, 4^. Despite extensive studies linking genetic impairments of mitochondrial homeostasis (e.g. altered fidelity of mtDNA synthesis, mitochondrial UPR, OXPHOS and others) to diseases and accelerated aging^2, 5, 6^, it is less clear what endogenous processes instigate mitochondrial decline during normal aging. The identification of such “natural” drivers of mitochondrial aging is however crucial because they likely comprise suitable intervention targets towards restoration of mitochondrial integrity and organismal health in late life. In this study, we combined omics, genetics and functional analyses in *C. elegans* and human to discover novel interventions that improve mitochondrial health and metabolic plasticity during advanced aging. Our tests revealed a decline in phosphatidylcholine synthesis as a novel conserved driver of natural mitochondrial aging, which can be overcome by dietary supplements.

## Main

### Proteomics analysis of wild type C. elegans and long-lived mitochondrial mutants reveals metabolic decline as a late event during aging

In our search for malleable processes that can alleviate mitochondrial deterioration with age, we decided to analyze genetic backgrounds that maintain normal or even extended longevity amid persistent mitochondrial dysfunction. While in most cases congenital mitochondrial impairments lead to cell dysfunction or accelerated aging^2^, there are two extensively characterized examples in *C. elegans* – the hypomorphic mutants *clk-1(qm30)* and *isp-1(qm150)*, which exhibit extended longevity despite carrying impaired mitochondria for the entire lifetime^7, 8^. It is thus feasible that these mutants possess intrinsic molecular adaptations, which support functionality of mildly impaired mitochondria. The previous omics studies comparing the mutants to WT counterparts were performed exclusively at young age^9–12^, identifying early life molecular differences without offering an aging perspective. To uncover potential adaptations with a life-long resolution, we performed label-free unbiased proteomics analysis in young (adulthood day 1, AD1), middle aged (adulthood day 5, AD5) and old (adulthood day 10, AD10) WT and mutant nematodes (Figure 1A) searching for protein expression changes, which occur in the aging WT but are alleviated in both mito-mutants. In this experiment, we measured a total number of 5339 proteins across all conditions (Table S1), and our proof of concept data analysis successfully detected previously described molecular features of the mito-mutants such as reduced ribosomal content in young *clk-1(qm30)* animals^13^ (Figure S1A and Table S3) and increased expression of the mitochondrial UPR components HSP-6 and HSP-60 in young *isp-1(qm150)* mutants^14^ (Figure S1B and Table S4). We next assessed protein changes occurring during early and advanced WT aging by analyzing proteins differentially regulated (Log2FC ≥ 0.58 and Q ≤ 0.05) between young (AD1) and post-reproductive (AD5), and young (AD1) and old (AD10) WT animals. The purpose of this experiment was to discover at what point mitochondria begin to fail during normal aging. We found a higher number of proteins (1835) to be differentially regulated in the AD10/AD1 comparison compared to the AD5/AD1 condition (1187) (Figure 1B and Tables S5-S7) in line with the expected stronger effect of aging in AD10 nematodes. We next extracted regulated proteins, which were either specific to early (AD5/AD1) or advanced (AD10/AD1) aging or overlapped between the two conditions (Table S7), and analyzed these by using the WormCat pathway enrichment analysis tool^15^. The analysis revealed alterations in mRNA splicing as the strongest enriched feature of early aging (Figure 1B and Table S8), while ribosomal changes and changes of stress responses were characteristic of both early and advanced aging (Figure 1B and Table S9). Finally, the advanced stage of aging was especially highly enriched in metabolic pathways including lipid and mitochondrial metabolism (Figure 1B and Table S10). This data suggested that alterations of proteostasis and stress responses occur relatively early during aging in line with previous reports^16, 17^, while metabolic and mitochondrial aging occur at a later point, consistent with our previous findings demonstrating functional loss of metabolic plasticity in nematodes on adulthood day 10^3^. Because our interest in studying mitochondrial mutants was mainly directed at the reversal of metabolic aging, we next compared protein sets differentially regulated between AD10 and AD1 in WT animals and the 2 mito-mutants. Initially, we observed that the number of proteins affected by aging in each strain was inversely correlated with reported lifespans of the animals: the highest number (1835) was seen in the shortest-lived WT strain (Figure 1C, E and Table S6) while the lowest number (1323) was found in the longest lived *isp-1(qm150)* mutant, with *clk-1(qm30)* nematodes residing in the middle both in terms of the number of affected proteins (1625) and the reported longevity^11^ (Figure 1C and E; Tables S11 and S15). This observation gave us additional reassurance of our data being representative of the aging processes in the strains. By combining WormCat analysis with the same data extraction strategy as in Figure 1B, we next determined that many terms indicative of metabolic aging showed similar enrichment patterns in WT and the two mito-mutants (Figure 1C-F and Tables S12-S18) despite the perceived metabolic resilience of the mutants. At the same time, we found distinct mitochondrial processes to be specifically enriched in the aging mito-mutants in line with (a) the mitochondrial origin of their differential longevity and (b) their functionally distinct mitochondrial impairments (Figure 1C and 1E; Tables S14 and S18). Finally and strikingly, when 2 mitochondrial mutants were compared together, only a small fraction of the overlap between their aging proteome profiles was assigned to mitochondria (Figure 1G and H, and Tables S19-S22), suggesting that the putative common adaptation to lifelong mitochondrial dysfunction might comprise non-mitochondrial processes. Of note, the terms highlighted by individual enrichment in the mutants such as chromatin and histone modifications and mitochondrial ribosome (Figure 1G, and Tables S21 and S22), were previously linked to both longevity extension and moderate mitochondrial dysfunction^18, 19^, confirming that the chosen data analysis approach delivers physiologically relevant outputs.

**Figure 1.**
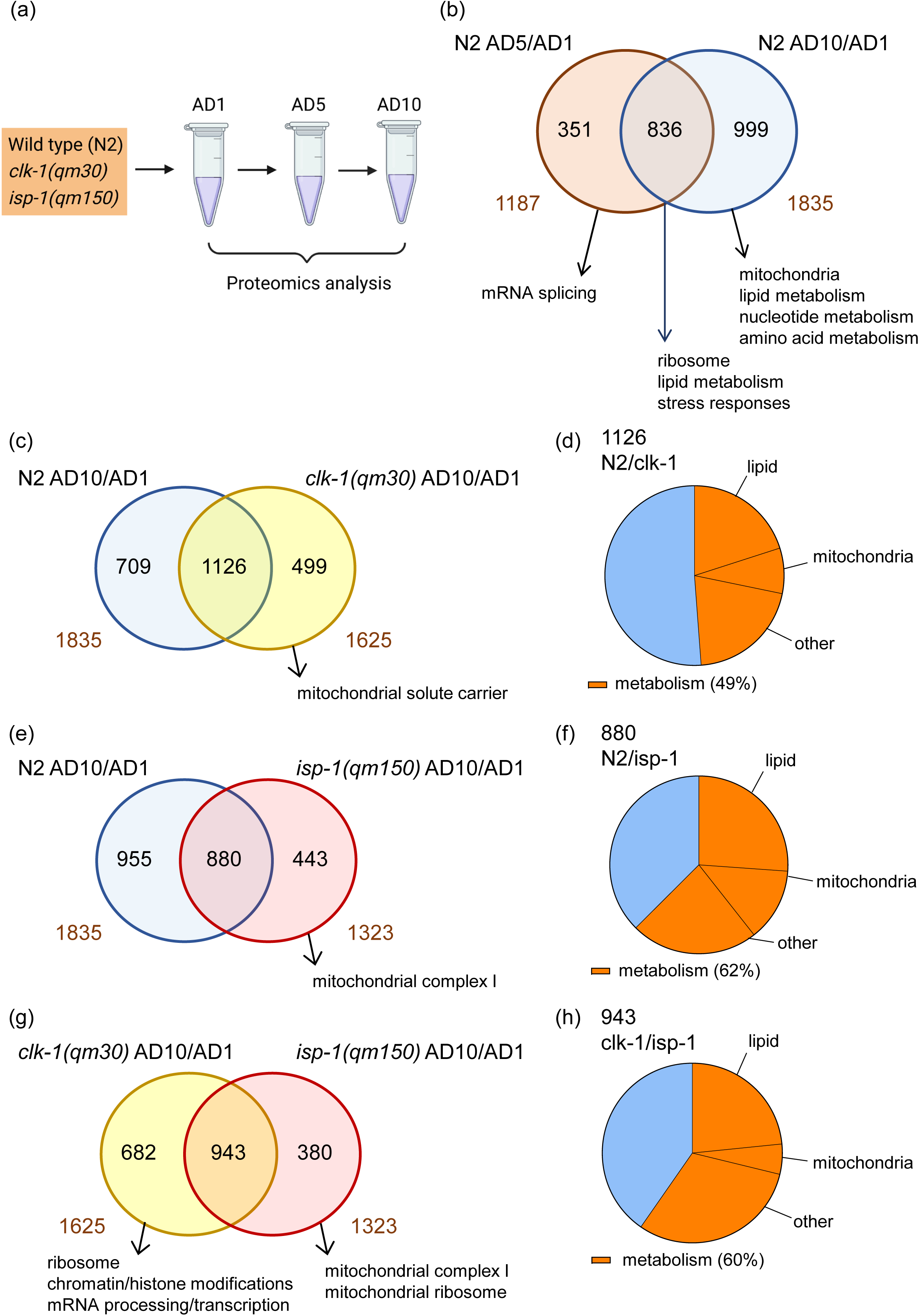
Metabolic alterations represent a late event during aging. Wild type (N2), *isp-1(qm150)* and *clk-1(qm30)* nematodes were age-synchronized, and proteomics samples (n ≥ 500 animals per sample) were collected at young (adulthood day 1, AD1), post-reproductive (AD5) and old (AD10) age **(a)**. **(b)** Differentially expressed proteins (DEPs, absolute Log2FC ≥ 0.58 and Q ≤ 0.05) in post-reproductive (AD5 vs AD1, N2 AD5/AD1) and old (AD10 vs AD1, N2 AD10/AD1) wild type animals were compared, revealing overlapping and non-overlapping subsets. Each of these subsets was analyzed using WormCat pathway analysis tool. Venn diagram and respective WormCat analysis highlights are shown. Numbers outside the diagram depict total DEPs in each condition, numbers inside the diagram represent respective overlapping and non-overlapping DEPs. Complete lists of DEPs used as an input, Venn diagram sub-setting and full WormCat outputs for **b** can be found in Tables S5-S10. **(c)** Proteins differentially regulated in old wild type animals (AD10 vs AD1, N2 AD10/AD1) and old *clk-1(qm30)* mutants (AD10 vs AD1, *clk-1(qm30)* AD10/AD1) were compared, analyzed and presented as in **b**. Respective protein lists and WormCat results, assembled as for panel **b**, are shown in Tables S11-S14. **(d)** Proteins differentially regulated in old wild type animals (N2 AD10/AD1) and old *isp-1(qm150)* mutants (*isp*-*1(qm150)* AD10/AD1) were compared, analyzed and presented as in **b**. Accessory lists and analyses are shown in Tables S15-S18. **(e)** Proteins differentially expressed in old *clk-1(qm30)* (*clk-1(qm30)* AD10/AD1) and *isp-1(qm150)* (*isp*-*1(qm150)* AD10/AD1) mutants were compared, analyzed and presented as in **b**. Accessory lists and analyses are shown in Tables S19-S22. **(d, f, h)** partitioning of overlapping DEPs from panels **c**, **e** and **g** respectively into metabolic and non-metabolic pathways based on the WormCat analysis is shown. Accessory WormCat-based calculations are presented in Tables S13, S17 and S20 respectively. Statistical analysis was adopted from WormCat.

### S-adenosylmethionine synthase SAMS-1 safeguards longevity in the context of mitochondrial impairments and is downregulated during normal aging

In order to continue probing the common “mitochondria-support” pathways between the 2 mito-mutants, we next searched for individual proteins whose expression is strongly and progressively affected by advanced aging in WT animals, with changes alleviated in the aging mito-mutants. We started by the unbiased filtering of all regulated proteins in the WT AD10/AD1 data set to select the strongest changes with AVG Log2FC ≥ 2,5 followed by their Q value ranking in the ascending order, placing most significant changes on top of the list (Table S23). We next prioritized for downregulated proteins, which might indicate loss of function with aging, and this analysis returned S-adenosylmethionine synthase SAMS-1 as a clear top candidate (Table S23). Further analysis showed that the reduction of SAMS-1 protein expression in WT animals was indeed progressive between AD5 and AD10 age points (Figure 2A and B, and Table S4), and this change was alleviated in both mito-mutant backgrounds and at both tested aging stages (Figure 2A and B, and Table S4). Moreover, one carbon metabolism (1CC) – the pathway, which involves *sams-1*^20^, was indeed among the top pathway enrichment outputs, which overlapped between the 2 aging mito-mutants (Table S20). Collectively, these data indicated that elevated SAMS-1 levels may contribute to lifespan extension in the context of mitochondrial impairments. To experimentally test this hypothesis, we next performed the RNAi-mediated knock down (KD) of the *sams-1* gene in WT and mito-mutant strains (Figure S1C). We found that *sams-1*KD elicits lifespan extension in WT animals (Figure 2C) in line with previous reports^21, 22^, but strikingly the lifespan of both long-lived mito-mutants was significantly reduced upon *sams-1* KD (Figure 2D and E). These findings confirmed that *sams-1* has a specific role in supporting homeostasis and longevity in the context of persistent mitochondrial impairments.

**Figure 2.**
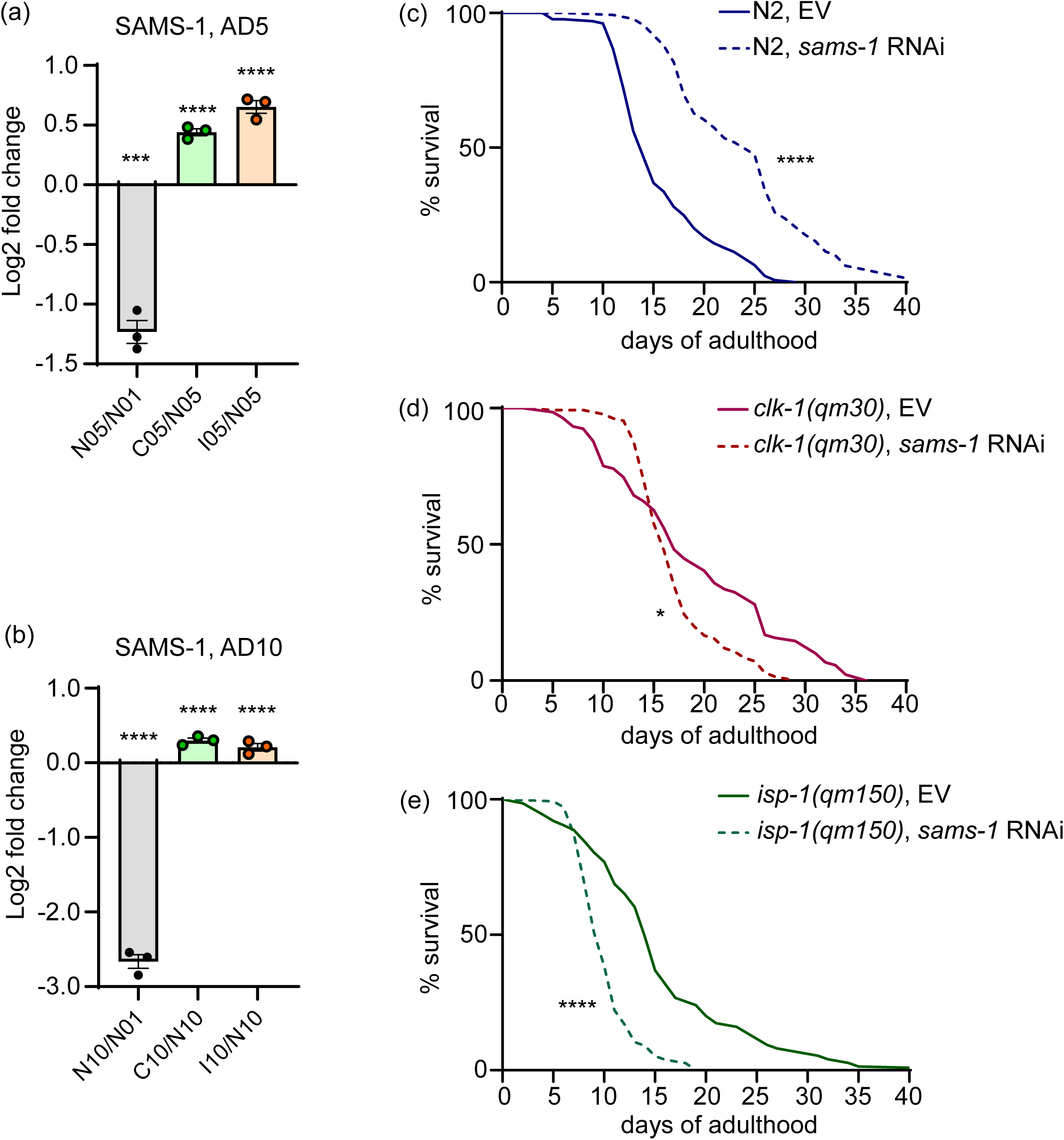
S-adenosylmethionine synthase SAMS-1 safeguards longevity in the context of mitochondrial impairments. Proteomics samples were collected as described in Figures 1a-b, and DEPs with strongest downregulation during wild type aging were identified as shown in Table S23. **(a)** The relative SAMS-1 levels (expressed as Log 2 fold change) in post-reproductive WT animals (N05/01, N2 AD5 vs AD1) as well as between post-reproductive *clk-1(qm30)* and WT (C05/N05, *clk-1(qm30)* AD5 vs N2 AD5) nematodes and *isp-1(qm150)* and WT (I05/N05, *isp-1(qm150)* AD5 vs N2 AD5) *C. elegans* are shown. **(b)** The same analysis as in was performed in old (AD10) animals, and relative SAMS-1 expression values are presented. In **a-b**, mean and SEM values are presented. **(c)** WT and mito-mutant animals were age synchronized and exposed to *sams-1* RNA from L1 larval stage, survival was scored daily, n=140 in each experimental condition. The accessory proteomics data for **a-b** can be found in Table S4. In **(a-b)** statistics was calculated using unpaired t-test and Tukey’s multiple comparison test, in **(c)** Mantel-Cox test was used, two-tailed *p* values were calculated. *-*p<0.05*; ***-*p<0.001* and ****-*p<0.0001*. Exact *p* values can be found in the statistics Source data file.

### sams-1 gene inactivation disrupts mitochondrial network integrity and triggers mitochondrial unfolded protein response

We next tested if *sams-1* gene inactivation has a direct impact on mitochondrial integrity and dynamics by subjecting the reporter strain harboring GFP labelled mitochondria in the body wall muscle^23, 24^ to *sams-1* RNAi. Strikingly, *sams-1* KD resulted in the severe disruption of the mitochondrial network as seen by loss of tubular mitochondria paralleled by an increase of fragmented and very fragmented organelles in the *sams-1* inactivated animals (Figure 3A and B). Because mitochondrial fusion is implicated in supporting mitochondrial efficacy and homeostatic integrity^25, 26^, we next assessed levels of mitochondrial stress by performing *sams-1* KD in the *C. elegans* strain expressing the mitochondrial unfolded protein response (UPR^mt^) reporter *hsp-6p::GFP*^14^. We found elevated levels of GFP expression in the reporter animals exposed to *sams-1* RNAi (Figure 3C and D) consistent with the requirement of *sams-1* for the stress-free mitochondrial homeostasis. Collectively, our data revealed a prominent role of *sams-1* in safeguarding mitochondrial integrity.

**Figure 3.**
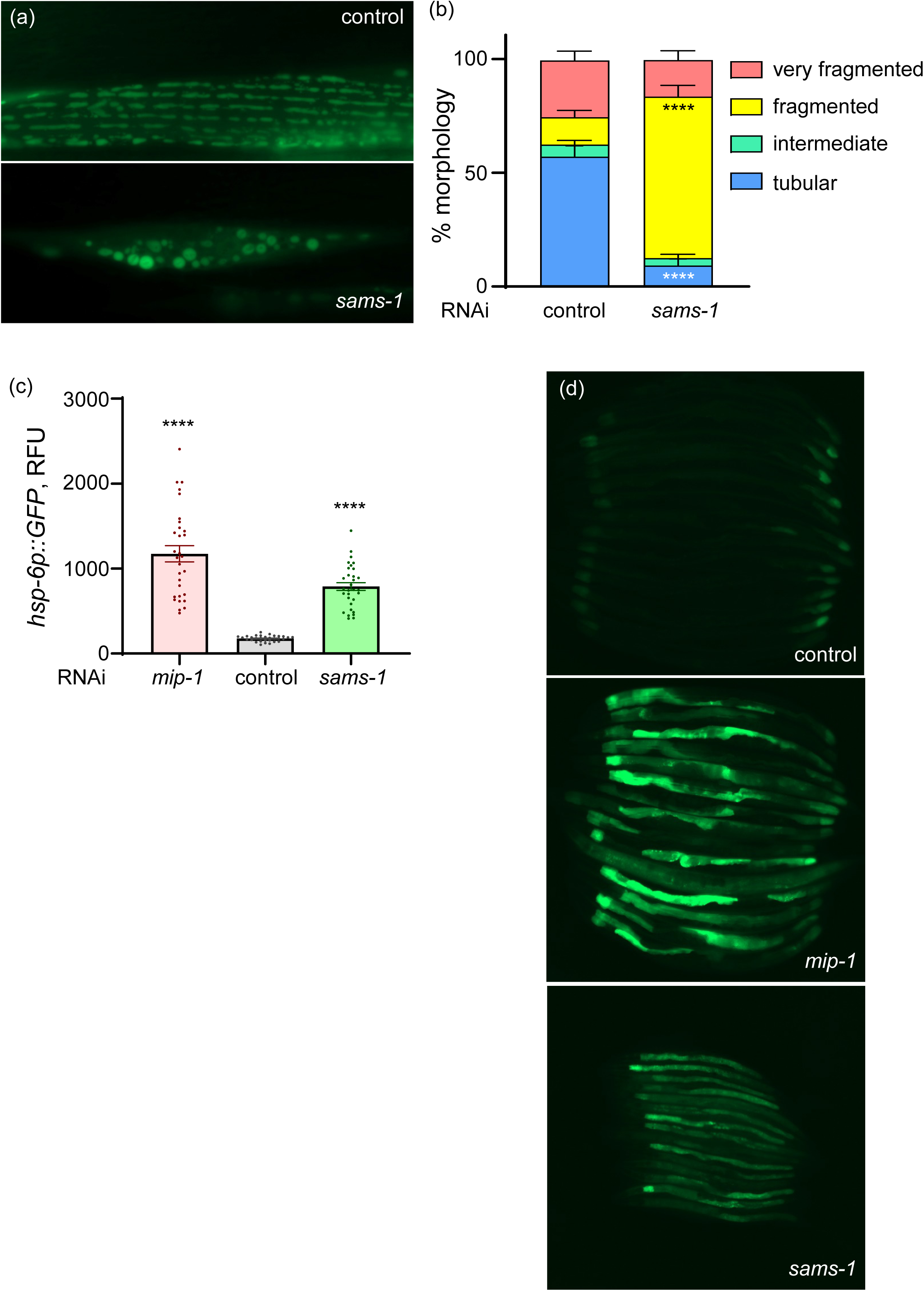
*sams-1* inactivation causes accelerated loss of mitochondrial integrity. **(a-b)** Transgenic animals expressing GFP-tagged mitochondria in the body wall muscle (*myo-3p::gfpmit*) were age-synchronized and exposed to *sams-1* RNAi or empty vector control from the L1 stage. Equally scaled, representative images of control and *sams-1* knock down (KD) mitochondria are shown in **(a)**. **(b)** The % of tubular, intermediate, fragmented, and very fragmented mitochondrial morphologies in *sams-1* KD and control nematodes were scored in young actively reproducing adults (day 2 of adulthood, AD2), n=60 in each condition. **(c-d)** Transgenic nematodes expressing GFP under control of the *hsp-6* mitochondrial chaperone gene promoter (*hsp-6p::GFP*) were age synchronized and exposed to control and *sams-1* RNAi from the L1 stage with *mip-1* RNAi serving as a positive control for the UPR MT induction. GFP expression was assessed microscopically and quantified on AD2, n≥30 in each condition **(c)** and equally scaled representative images are shown in **(d)**. Statistics in **b** and **c** was assessed using unpaired t-test with Welch’s correction, mean and SEM values are presented; two-tailed *p* values were calculated. ****-*p<0.0001*. Representative results of at least 3 independent experiments are shown. Exact *p* values can be found in the statistics Source data file.

### sams-1 and S-adenosylmethionine influence mitochondria through their role in the synthesis of phosphatidylcholine

The *sams-1* gene encodes the *C. elegans* orthologue of the S-adenosylmethionine synthase, which is responsible for the production of the methyl donor S-adenosylmethionine (SAM)^20^ (Figure 4A). We next asked if SAM depletion was causal of mitochondrial distortions induced by the *sams-1* KD (Figure 3). In line with this hypothesis, the dietary SAM supplementation alleviated mitochondrial fragmentation and dampened mitochondrial UPR activity in *sams-1* RNAi exposed nematodes (Figure 4B-D), clearly confirming the key role of SAM synthesis in mitochondrial protection by *sams-1*. As indicated previously, SAM is the key methyl donor for the variety of cellular processes including DNA and histone methylation as well as synthesis of the membrane lipid phosphatidylcholine (PC)^27^ (Figure 5A). Although the methylation-dependent pathway represents only one of the 2 avenues of cellular PC synthesis^28^ (Figure S2A), previous studies reported a significant and physiologically meaningful reduction of PC levels in the *sams-1* deficient nematodes^29^. Because the lipid composition of the mitochondrial membranes plays a role in mitochondrial fission and fusion^30^, and PC is the most abundant lipid in the mitochondrial membranes^31^, we next asked if mitochondrial phenotypes seen in the *sams-1* KD animals were driven by the disruption of PC synthesis. In the methylation-dependent branch of *C. elegans* PC synthesis, SAM is processed by phosphoethanolamine N-methyltransferases *pmt-1* and *pmt-2*^32, 33^ (Figures 5A and S2A). We next monitored mitochondrial morphology and UPR^mt^ induction in nematodes exposed to *pmt-1* RNAi (Figure 5E) and found that *pmt-1* inactivation replicated loss of mitochondrial integrity caused by the *sams-1* KD (Figure 5B), while only a marginal effect of *pmt-1* RNAi on the *hsp-6p::GFP* expression was detected (Figure 5C and D). These data suggested that a reduction of PC synthesis is likely responsible for the mitochondrial integrity/fusion defects elicited by the *sams-1* deficiency, while the strong enhancement of UPR^mt^ activity in *sams-1* KD animals is driven by a PC-independent mechanism. Interestingly, previous reports linked elevated UPR^mt^ activity to a reduction in histone methylation^18^, and similar decline of methylation could arise from the depletion of SAM in *sams-1* KD animals^34^ thus inducing UPR^mt^ by the chromatin-dependent non-mitochondrial mechanism. Accordingly, the SAM-dependent re-installment of histone methylation could explain the dampening of the UPR^mt^ reporter expression in *sams-1* deficient animals supplemented with SAM (Figure 4C and D).

**Figure 4.**
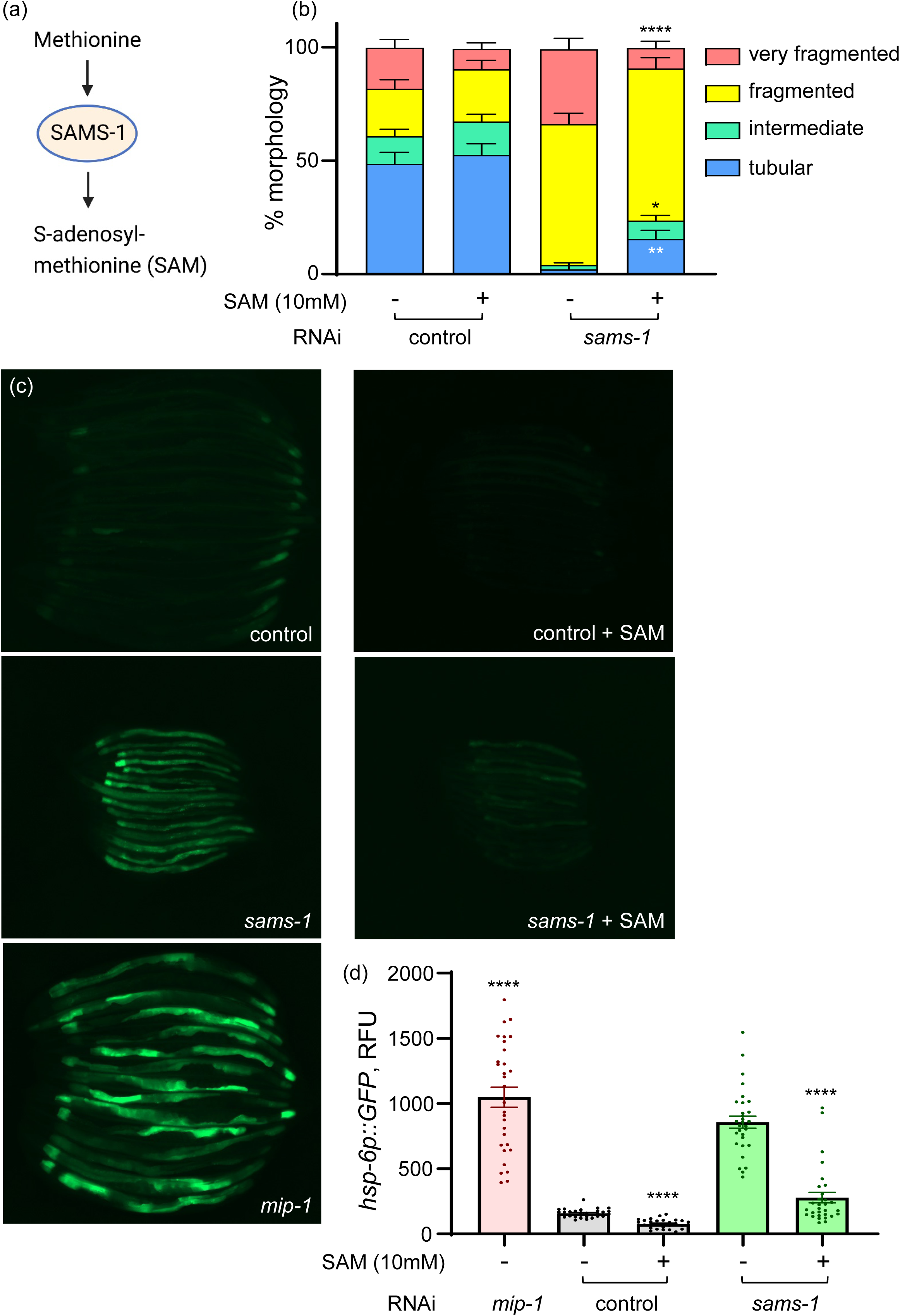
SAMS-1 supports mitochondrial integrity via S-adenosylmethionine synthesis. **(a)** Schematic depicts the role of SAMS-1 in converting methionine to S-adenosylmethionine (SAM). **(b)** Transgenic nematodes expressing GFP-tagged mitochondria in the body wall muscle (*myo-3p::gfpmit*) were age-synchronized and exposed to *sams-1* RNAi or empty vector control from the L1 stage, 10mM SAM was supplemented from the L4 stage. Mitochondrial morphologies were scored on AD2 as in Figure 3b, n=60 in each experimental group. **(c-d)** Transgenic nematodes expressing GFP under control of the *hsp-6* promoter (*hsp-6p::GFP*) were age synchronized and exposed to control and *sams-1* RNAi from the L1 stage, 10mM SAM was given from the L4 stage; *mip-1* RNAi was used as positive control for the UPR MT induction. GFP expression was assessed microscopically and quantified on AD2, n=30 in each condition **(d)** and equally scaled representative images are shown in **(c)**. Statistics in **b** and **d** was assessed using unpaired t-test with Welch’s correction, mean and SEM values are presented; two-tailed *p* values were calculated. *-*p*<0.05; **-*p*<0.01; ****-*p<*0.0001. Representative results of at least 3 independent experiments are shown. Exact *p* values can be found in the statistics Source data file.

**Figure 5.**
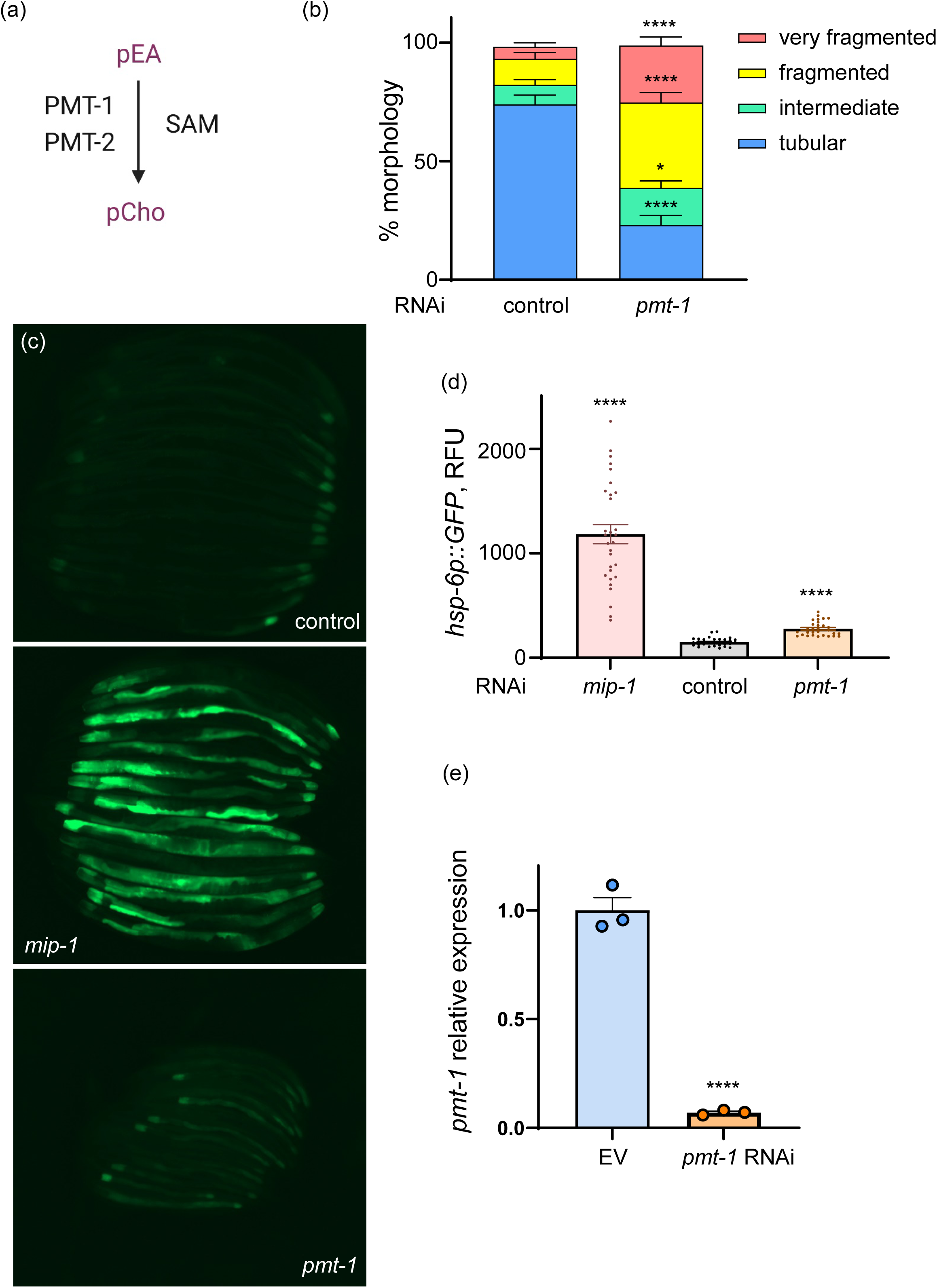
SAM-dependent phosphatidylcholine synthesis links *sams-1* to modulation of mitochondrial morphology. **(a)** Schematic depicts the pathway of converting phosphatidylethanolamine (pEA) to phosphatidylcholine (pCho) in a SAM-dependent manner by phosphoethanolamine methyltransferases 1 and 2 (PMT-1 and 2). **(b)** Transgenic nematodes expressing GFP-tagged mitochondria in the body wall muscle (*myo-3p::gfpmit*) were age-synchronized and exposed to *pmt-1* RNAi or empty vector control from the L1 stage. Mitochondrial morphologies were scored on AD2 as in Figure 3b, n=60 in each experimental group. **(c-d)** Transgenic nematodes expressing GFP under control of the *hsp-6* promoter (*hsp-6p::GFP*) were age synchronized and exposed to control and *pmt-1* RNAi from the L1 stage; *mip-1* RNAi was used as positive control for the UPR MT induction. GFP expression was assessed microscopically and quantified on AD2, n=30 in each condition **(d)** and equally scaled representative images are shown in **(c)**. **(e)** The result of quantitative RT PCR depicting *pmt-1* mRNA expression in control and *pmt-1* RNAi treated worms is shown. Statistics in **b** and **d** was assessed using unpaired t-test with Welch’s correction and in **e** – by unpaired t-test, mean and SEM values are presented; two-tailed *p* values were calculated. *-*p*<0.05; ****-*p<*0.0001. Representative results of at least 3 independent experiments are shown. Exact *p* values can be found in the statistics Source data file.

We next tested the effect of the dietary PC (pCho) supplementation on the mitochondrial and organismal phenotypes elicited by *sams-1* and *pmt-1* KDs. In line with PC having no impact on the UPR^mt^ activity in *sams-1* inactivated nematodes, PC supplementation failed to dampen the *hsp-6p::GFP* expression in these KD animals (Figure S2B and C). At the same time, PC supplementation alleviated the excessive mitochondrial fragmentation (Figure 6A and B) and the reduction of body size (Figure 6C) in both *sams-1* and *pmt-1* KD worms. The body size alteration could also be rescued by choline (Cho) supplementation (Figure 6D). Choline is converted to PC by the conserved choline kinase (CK)/CPD-choline pathway that is complementary to the methylation-dependent PC synthesis (Figure S2A) and is active in most tissues^28, 35^. The ability of choline supplementation to rescue the size defects of *sams-1* and *pmt-1* KD worms is thus supportive of the link between PC levels and organismal and mitochondrial fitness. Of note, reduction of body size was previously linked to lowered PC synthesis in nematodes^29^ but no connection to mitochondrial integrity was previously made.

**Figure 6.**
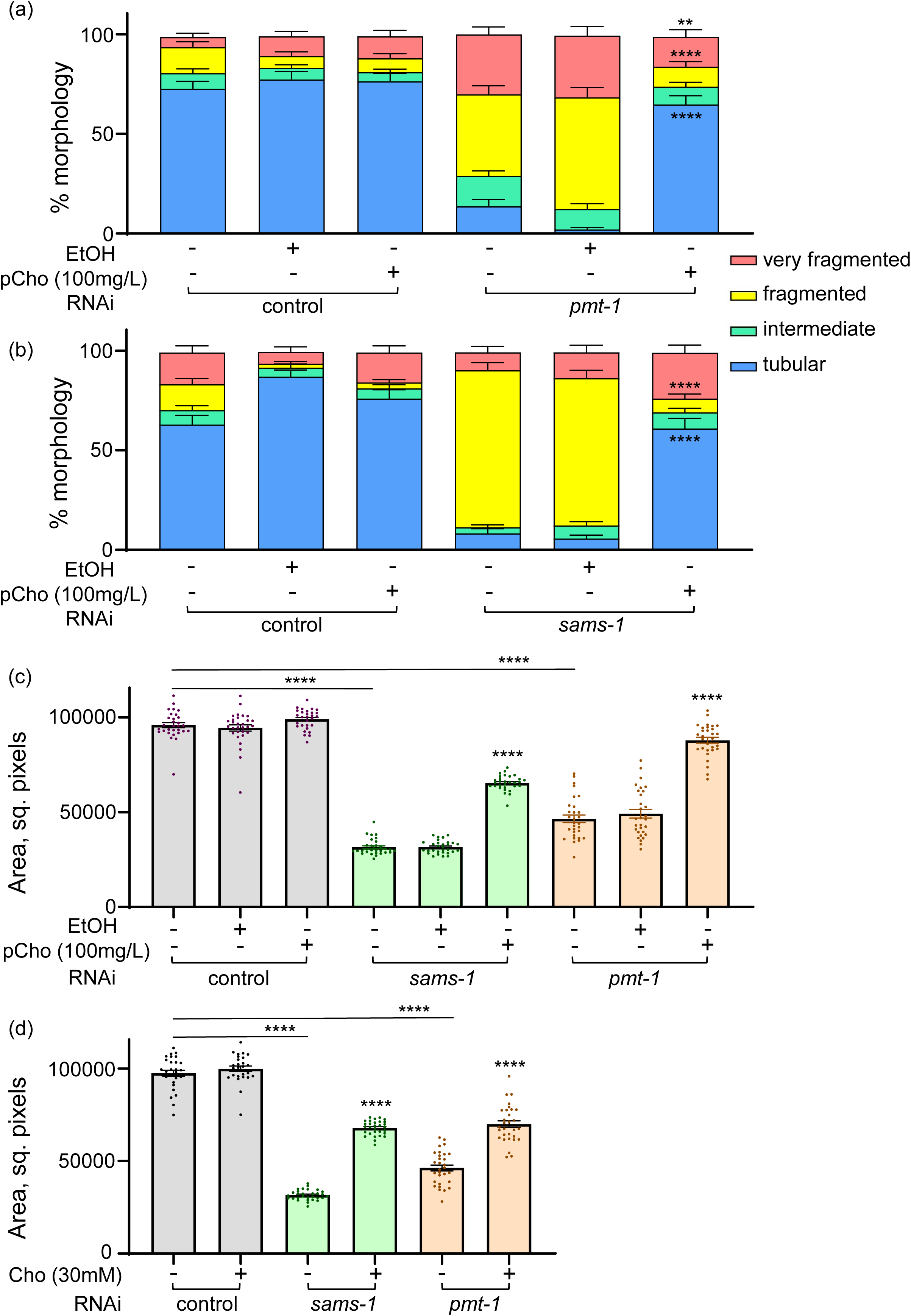
Phosphatidylcholine feeding rescues mitochondrial morphology and body size defects of *sams-1* and *pmt-1* deficient nematodes. **(a-b)** Transgenic nematodes expressing GFP-tagged mitochondria in the body wall muscle (*myo-3p::gfpmit*) were age-synchronized and exposed to *pmt-1* RNAi **(a)** or *sams-1* RNAi **(b)** with empty vector RNAi as control from the L1 stage, 100mg/L phosphatidylcholine (pCho) was provided from L4 stage and 0,1% EtOH was used as a vehicle control for pCho. Mitochondrial morphologies were scored on AD2 as in Figure 3b, n=60 in all experimental groups. **(c)** Age-synchronized nematodes were exposed to RNAi and compound treatment as in **(a-b)** and body size (area) was measured on AD2, n≥29 in each condition. **(d)** Age-synchronized nematodes were exposed to RNAi as in **(a-b)** and 30mM choline (Cho) was given from L4 stage, body size (area) was measured on AD2, n≥28 in each condition. Statistics in **a-d** was assessed using unpaired t-test with Welch’s correction, mean and SEM values are presented; two-tailed *p* values were computed. ****-p<0.0001. Representative results of at least 3 independent experiments are shown. Exact *p* values can be found in the statistics Source data file.

### Ectopic PC supplementation alleviates mitochondrial aging and restores metabolic resilience under stress

We next asked if the reduction of methylation-dependent PC synthesis was a relevant phenomenon in the context of normal aging possibly contributing to aging-associated mitochondrial and cellular dysfunction^3^. We first re-examined the most outstanding protein changes occurring in aging WT nematodes and notably found PMT-1, PMT-2 and SAMS-1 to be the top 3 strongest downregulated proteins at the advanced age (AD10) (Table S23). Moreover and similar to changes shown for SAMS-1 in Figure 2A and 2B, the downregulation of PMT-1 and PMT-2 expression was discovered to be progressive with age (Figure 7A and S3A, Table S4). These findings highlighted methylation-dependent PC synthesis as one of the strongest aging-inhibited pathways in *C. elegans*. We next tested if the expression of PEMT - a functional orthologue of PMT-1/2 in humans, also declines during aging by analyzing the transcriptomic data of the Genotype-Tissue Expression (GTEx) project^36, 37^. Interestingly, a trend of PEMT down-regulation with age was observed across several human organs (Figure S4 and Table S26) with highest PEMT expressing tissues (top 25%) showing the most notable reduction (Figure 7B, Figure S4, Tables S26 and S27). Of note, the especially affected cohort included organs with high lipid content such as subcutaneous and visceral adipose tissues (Figure 7C and S3C; Tables S28 and S30) and the ovary (Figure S3B and Table S29). These observations indicated that the decline of methylation-dependent PC synthesis is occurring also in human aging at least in a subset of relevant cell types.

**Figure 7.**
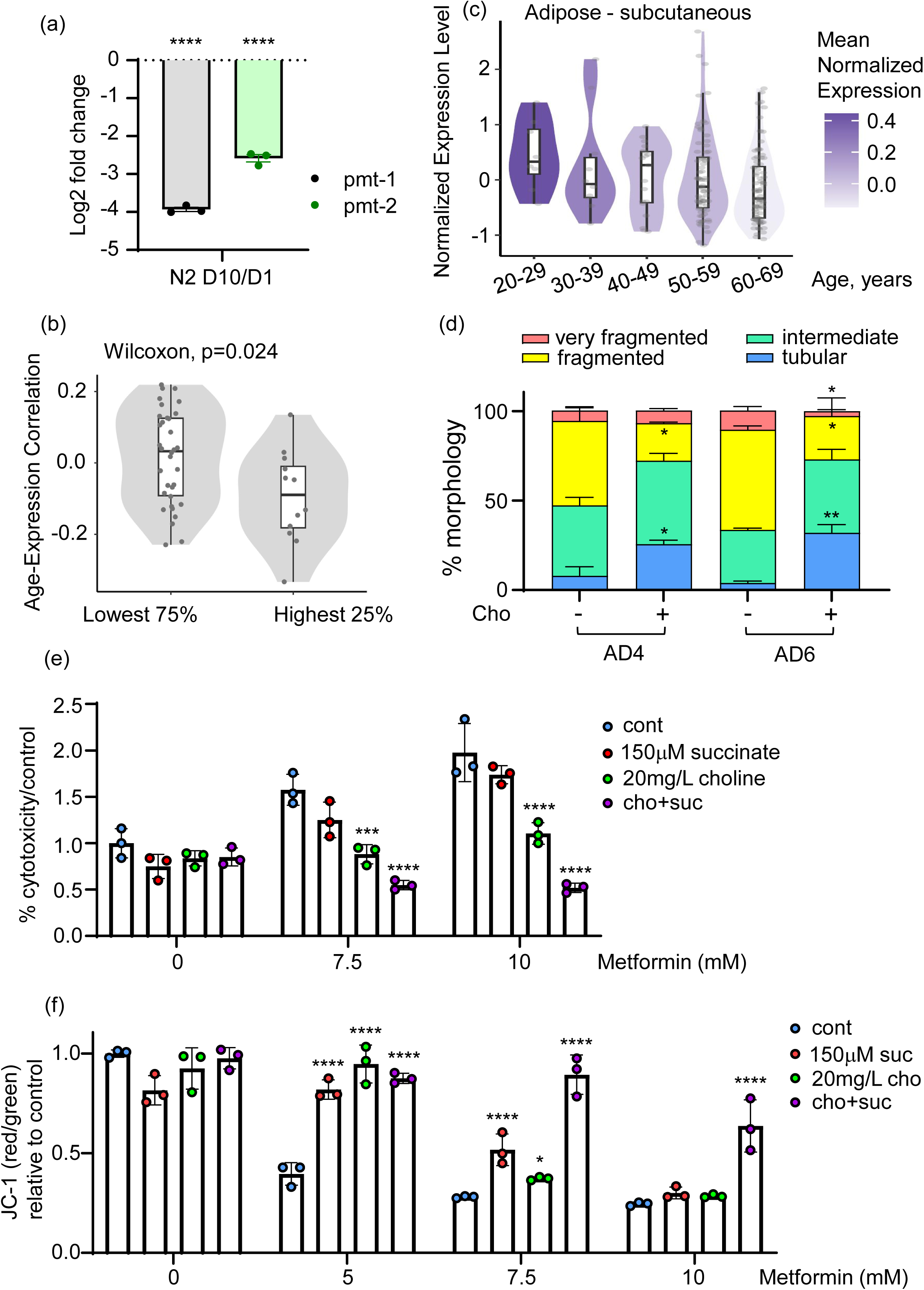
A decline of phosphatidylcholine synthesis is a malleable inducer of mitochondrial dysfunction during aging. **(a)** Proteomics samples were collected as described in Figures 1a-b and relative levels (expressed as Log 2 fold change) of PMT-1 and PMT-2 proteins in old (AD10 vs AD1) WT animals are shown. All relevant calculations can be found in Table S4. **(b)** The expression of PEMT gene across human tissues at different age was analyzed using the GTEx dataset (v8). The selection of samples, details of data processing and exclusion criteria can be found in the methods section. Tissues with high (highest 25%) and low (lowest 75%) PEMT expression were determined as described in Figure S4, and age-related changes in each tissue were computed using Spearman’s correlation between age and expression level across all available age points. The comparison of age-related changes between PEMT high- and low-expressing tissues is shown, and complete data can be found in Table S27. **(c)** The expression of human PEMT gene in the subcutaneous adipose tissue at different age was tested using the GTEx dataset (v8) as described in (b). Normalized expression levels represent individual log2 transformed quantile normalized Transcripts Per Million (TPM) values corrected for sex and death circumstances using a linear model; median normalized PEMT expression in each age group is color coded and complete data can be found in Table S28. **(d)** Transgenic nematodes expressing GFP-tagged mitochondria in the body wall muscle (*myo-3p::gfpmit*) were age-synchronized and grown with and without 30mM Cho supplementation for indicated times. Mitochondrial morphologies were scored on AD4 and AD6 as described in Figure 3b, n≥54 in each experimental group. **(e-f)** BJ human skin fibroblasts were treated with indicated concentrations of metformin with or without 150μM succinate, 20mg/L choline or a mix of choline and succinate. Cytotoxicity (LDH assay, **e**) and mitochondrial membrane potential (JC-1 assay, **f**) were measured after 24h. Significance was assessed by unpaired t-test in **a** and **d,** Wilcoxon test in **b** and multiple comparison t-test in **e-f**, mean and SEM values are presented; two-tailed *p* values were computed. *-*p*<0.05; **-*p*<0.01; ***-*p*<0.001; ****-*p<*0.0001. Representative results of at least 3 independent experiments are shown. Exact *p* values can be found in the statistics Source data file.

A reduction of SAM-dependent PC synthesis was previously linked to pathological elevation of triglyceride (TAG) storage in the rat liver^38^, and increased abundance/size of TAG containing lipid droplets was seen in *sams-1* and *pmt-1* deficient nematodes^29, 34, 39^. In earlier work, we showed that lipid droplet abundance can be probed by measuring the expression of their core protein components vitellogenins^3^, and indeed vitellogenin expression was progressively elevated with age in WT *C. elegans* (Figure S5A and Table S24). This finding coincided with aging-triggered decline of SAMS-1, PMT-1 and PMT-2 protein levels (Figures 2A-B, 7A and S3A), suggesting that reduced PC synthesis may contribute to the de-regulation of organismal lipid storage also in the context of aging. Previous tests in young nematodes implicated increased UPR^ER^ activity in connecting lowered PC levels to the elevated storage of TAGs, demonstrating increased expression of ER chaperones HSP-3 and HSP-4 in *pmt-2* deficient *C. elegans*^39^. However, we found HSP-3 and HSP-4 levels to be reduced with age in WT nematodes (Figure S5B and C; Table S4) suggesting that UPR^ER^ has no role in connecting lowered PC synthesis with elevated lipid droplet abundance during aging. On the other hand, our earlier work implicated aging-associated mitochondrial dysfunction in driving the TAG storage abnormalities in late life^3^. We thus hypothesized that in old cells the reduction of PC synthesis may cause alterations of lipid turnover via interference with mitochondrial integrity. Notably, the increased mitochondrial fragmentation and reduced network integrity seen in both *sams-1* and *pmt-1* KDs (Figures 3-5), are well known features of aged mitochondria^3, 40^. In the next experiment we on the one hand validated the previously described increase of mitochondrial fragmentation during normal aging^3, 40^, and on the other hand determined that dietary choline supplementation indeed alleviates this aging-triggered alteration (Figure 7D). We used choline instead of PC in this experiment to avoid long-term exposure of the animals to organic solvents required for the PC supplementation in the context of NGM agar, and because choline kinase components mediating the conversion of choline to PC did not show a coordinated decline in expression during aging (Figure S3D and Table S25). Collectively, our data demonstrate that a reduction of methylation-dependent PC synthesis contributes to loss of mitochondrial integrity during normal aging with possible broader effects on cellular lipid storage. We also demonstrate that the aging-linked alteration of mitochondrial fusion can be alleviated by dietary boosting of choline/PC levels.

Mitochondrial integrity and specifically mitochondrial fusion are important regulators of metabolic plasticity that are known to decline with age affecting cellular health and responses to drugs like metformin^2, 3^. Specifically, a reduction of mito-fusion contributes to metformin toxicity in old nematodes and cells^3^. We next tested the effect of choline supplementation in the human cell culture model of metformin toxicity^3^ in comparison or combination with the electron transport chain Complex II substrate succinate. The boosting of Complex II activity was chosen as it compensates for the inhibition of Complex I by metformin^41^. We found that choline protected metformin-exposed cells from cell death and loss of mitochondrial membrane potential similar to succinate, with most potent rescues delivered by a combination of both supplements (Figure 7E and F). These data demonstrate that restoration of mitochondrial network integrity by PC repletion is protective against aging-induced mitochondrial fragmentation and has a restorative impact on mitochondrial plasticity across species.

## Discussion

In this study, we explored natural mechanisms that contribute to the deterioration of mitochondria during normal aging in order to identify targets mendable by pharmacological or dietary interventions. While congenital failures of mitochondria are widely known to cause premature aging^2, 5, 6^, the “natural causes” of mitochondrial aging are surprisingly less clear. We initially used longitudinal proteomics analysis in WT *C. elegans* and long-lived mitochondrial mutants, which was coupled to RNAi-mediated gene inactivation and longevity testing, to demonstrate that S-adenosylmethionine synthase SAMS-1 is required for longevity maintenance in the context of mitochondrial impairments. Specifically, we could confirm previous reports of increased lifespan of WT animals exposed to *sams-1* RNAi^21, 22^, while the same RNAi exposure was detrimental for the longevity of long-lived *clk-1(qm30)* and *isp-1(qm150)* strains carrying hypomorphic mitochondrial mutations. At the same time, we found SAMS-1 to be among the strongest downregulated proteins in old WT nematodes, and same strong and progressive downregulation with age was discovered for phosphoethanolamine N-methyltransferases PMT-1 and PMT-2, which utilize S-adenosylmethionine (SAM, the product of SAMS-1) for mediating the conversion of phosphatidylethanolamine (PE) to phosphatidylcholine (PC). We next explored the mechanistic basis of the longevity link between *sams-1* gene inactivation and mitochondrial impairments, and discovered that knock downs (KDs) of both *sams-1* and *pmt-1* genes cause an early life increase of mitochondrial fragmentation that is comparable to mito-morphology alterations observed during normal aging^3, 40^. Importantly we could alleviate these morphology defects by dietary provision of PC^42^ or choline (can be converted to PC by the highly conserved CDP-choline pathway^35^) both in the KDs and also during WT aging. Finally, we discovered by using the GTEx dataset (v8)^36^ and previously pre-processed gene expression data^37^ that levels of the human PMT-1/2 analogue PEMT decline with age in several tissues and especially in organs showing overall highest PEMT expression. This data indicates that aging-associated decrease of methylation-dependent PC synthesis is evolutionary conserved and may contribute to “natural” aging of mitochondria across species. Subsequently, we used human cell culture model of metformin toxicity to demonstrate that choline/PC repletion restores metabolic plasticity in human cells affected by mitochondrial stress.

In this study focusing on mechanisms of aging, we did not scrutinize the exact biochemical links between PC levels and mitochondrial morphology. PC is one of the most abundant lipids in the mitochondrial membrane^31, 43^ and a precursor of phosphatidic acid (PA)^44^. PA in turn is known to promote mitochondrial fusion both at the level of mitochondrial membrane structure (by creating negative membrane curvature) and by interacting with the fission and fusion machinery (PA interferes with Drp1-medated fission and stimulates the activity of mitofusins-1 and -2)^45, 46^. In addition, previous reports found significant alterations of mitochondrial morphology and ultrastructure in cells lacking the PC precursor PE^43, 47^ (pEA in Figure S2A). These earlier findings provide the appropriate mechanistic context for the hereby discovered role of PC synthesis in the decline of mitochondrial plasticity during aging (Figure S6). In summary, we used a combination of omics and functional tests in two model systems (*C. elegans* and human) to identify aging-associated decline of methylation-dependent PC synthesis as a “natural” trigger of mitochondrial aging that is malleable by dietary treatments. Of note, both choline and PC are bioavailable in humans comprising supplements of clinical relevance^48, 49^.

Finally, the comparison of our data with the existing literature indicates that physiological stressors may be mitigated by diverse homeostatic pathways at different age. For instance, omics analyses of the same *C. elegans* strains as used in our proteomics study were already conducted earlier with the exclusive focus on young animals^9–12^. These earlier tests revealed increased activity of adaptive stress responses such as autophagy^12^ and DAF-16/FOXO targets^11^ as key longevity-relevant differences between WT nematodes and long-lived mito-mutants. The results of our longitudinal proteomics study agree with these findings in identifying a decline of proteostasis and stress responses as an early event during WT aging, and it is feasible that higher stress resilience confers a delayed onset of aging in the mito-mutants. However, as seen in our omics and functional data, when metabolic failures onset at the advanced age, their mitigation requires other protective mechanisms such as modulation of lipid synthesis and mito-morphology. Specifically, we hereby reveal mitochondrial fusion as a suitable and malleable target of anti-aging treatments focused on elderly patients.

## Supporting information

Statistics source data

## Acknowledgements

We thank the Proteomics Core Facility and especially Dr. Joanna Kirkpatrick, and SPARK Technology Transfer Core Facility at FLI for supporting this study. The FLI is a member of the Leibniz Association and is financially supported by the Federal Government of Germany and the State of Thuringia. TP was supported by the German Academic Exchange Services (Deutsche Akademische Austauschdienst, DAAD). ME and LE were supported by the EU-ESF Thüringer Aufbaubank funding (2019 FGR 0082). ME is funded by the ERC CoG LifeLongFit of the European Commission. ME is also funded by the Carl-Zeiss-Stiftung via IMPULS consortium and is a member of Cluster of Excellence Balance of the Microverse funded by the Deutsche Forschungsgemeinschaft (DFG). PA was supported by the DFG-funded Cluster of Excellence Balance of the Microverse. MD is funded by Carl-Zeiss-Stiftung (P2021-00-007). *C. elegans* strains were obtained from the Caenorhabditis Genetics Center (CGC), which is funded by NIH.

## Author contribution

ME conceptualized and designed the study; TP, PA, LE, MB and YL performed experiments; ME and TP analyzed the data; MD designed and performed human gene expression analysis; ME, TP and MD prepared figures; ME, TP and MD performed statistical analysis; ME wrote the manuscript and TP and MD reviewed the manuscript.

## Data and code availability

The mass spectrometry proteomics data have been deposited to the ProteomeXchange Consortium (Deutsch et al., 2020) via the PRIDE (Perez-Riverol et al., 2019) partner repository available at http://www.ebi.ac.uk/pride, with the dataset identifier PXD024180. Code for PEMT analysis: https://github.com/mdonertas/pc_synthesis_human_aging. Code for GTEx data preprocessing: https://github.com/mdonertas/aging_in_GTEx_v8.

---

**Figure S1.**
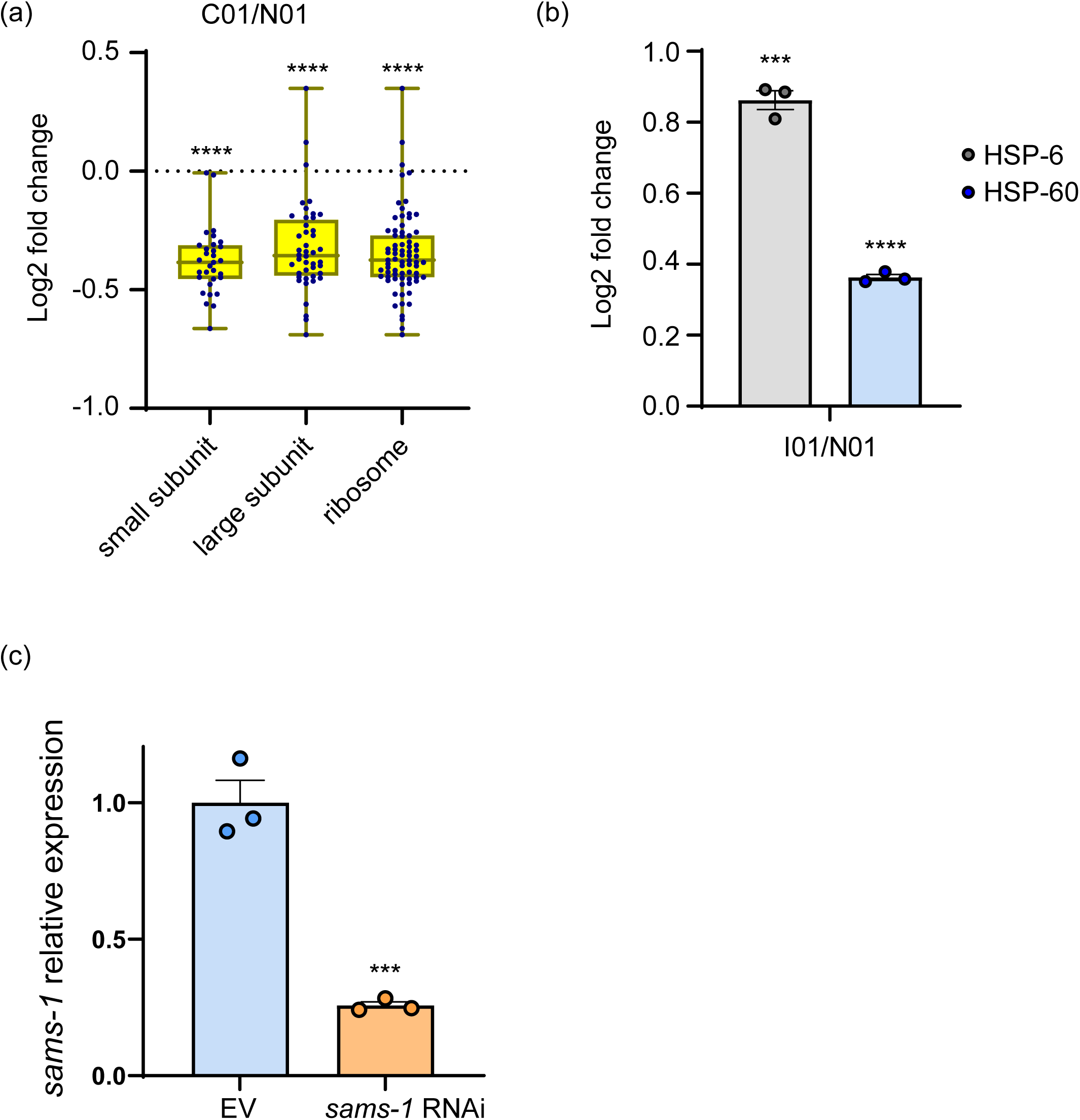
Validation of proteomics results and *sams-1* gene knock down efficiency. **(a-b)** Proteomics samples were collected as described in Figures 1a-b. Box plots depicting relative expression (Log 2 fold change) of all detected ribosomal proteins belonging to the small subunit, large subunit and whole ribosome between young *clk-1(qm30)* and WT nematodes are shown in **a**. Each dot represents one protein, median expression values are shown inside each box and whiskers highlight minimal and maximal values within each tested group. Statistics was assessed by Wilcoxon rank-sum test and all relevant expression values and calculations can be found in Table S3. **(b)** Relative expression values (Log 2 fold change) of HSP-6 and HSP-60 mitochondrial chaperones between young *isp-1(qm150)* and WT nematodes are presented. Each dot represents one replica sample, mean and SEM values are presented and unpaired t-test was used to compute *p*-values. All relevant expression values and calculations can be found in Table S4. **(c)** Expression of the *sams-1* gene was measured by quantitative RT-PCR in animals exposed to EV control RNAi and RNAi targeting *sams-1*. Result is representative of 3 independent tests, mean and SEM values are presented, unpaired t-test was used for the statistical assessment. ***- *p*<0.001; ****-*p*<0.0001, all values are two-tailed. Exact *p* values can be found in the statistics Source data file.

**Figure S2.**
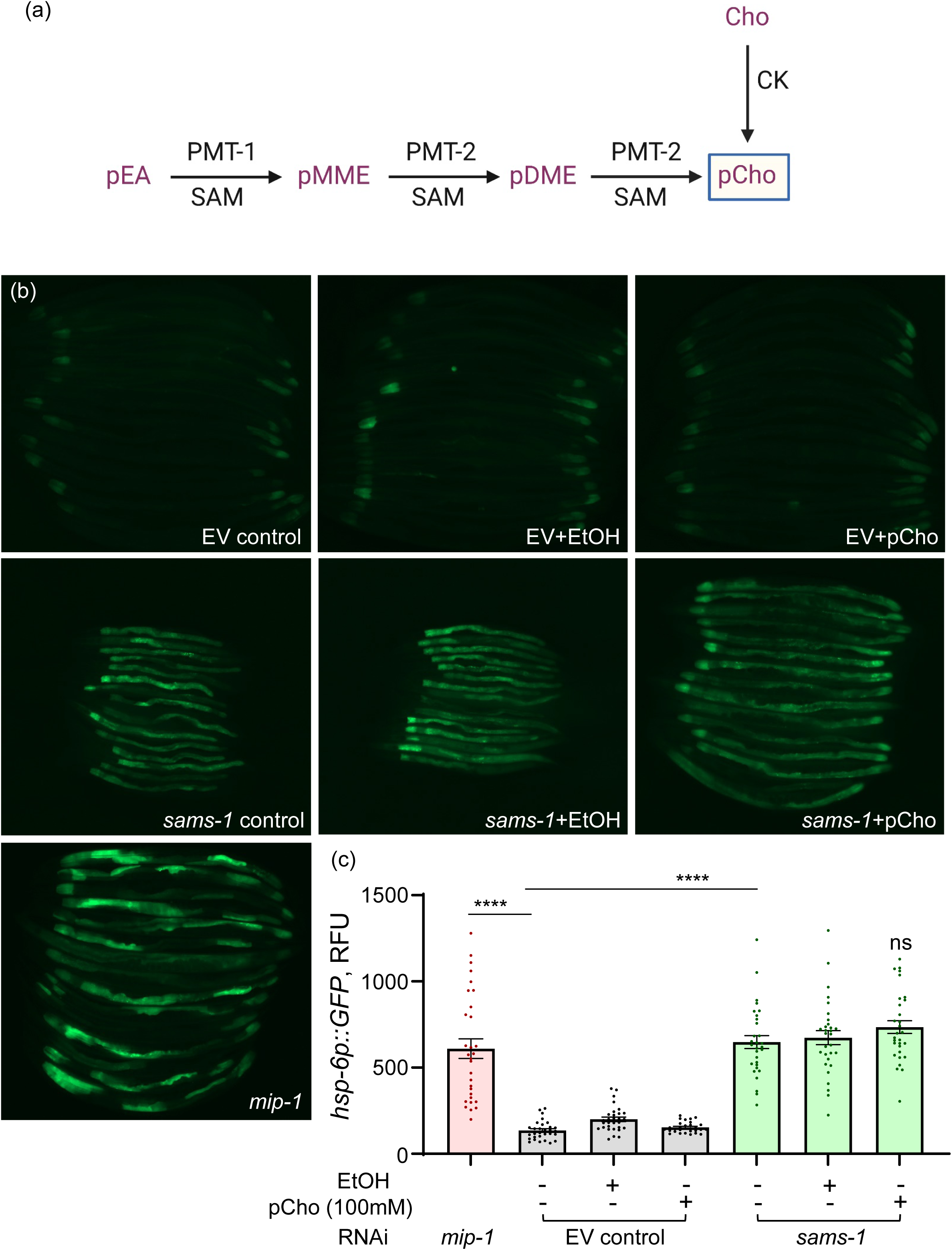
Phosphatidylcholine repletion doesn’t dampen high UPR^mt^ activity in *sams-1* knock down animals. **(a)** Schematic representation of SAM-dependent and independent pathways of Phosphatidylcholine (pCho) synthesis is shown. CK - choline kinase; pEA – phosphatidylethanolamine, pMME – phosphatidylmonomethylethanolamine; pDME – phosphatidyldimethylethanolamine, Cho – choline. **(b-c)** Transgenic nematodes expressing GFP under control of the *hsp-6* promoter (*hsp-6p::GFP*) were age synchronized and exposed to control and *sams-1* RNAi from the L1 stage, 100mg/L phosphatidylcholine (pCho) was provided from the L4 stage and 0,1% EtOH was used as a vehicle control for pCho; *mip-1* RNAi was used as positive control for the UPR MT induction. GFP expression was assessed microscopically and quantified on AD2, n≥29 in each condition. Equally scaled representative images are shown in **b** and quantification in **c**. Mean and SEM values are presented, statistics was assessed by using unpaired t-test with Welch’s correction and two-tailed *p*-values were computed. Results are representative of 3 independent tests. ****-*p*<0.0001, exact *p* values can be found in the statistics Source data file.

**Figure S3.**
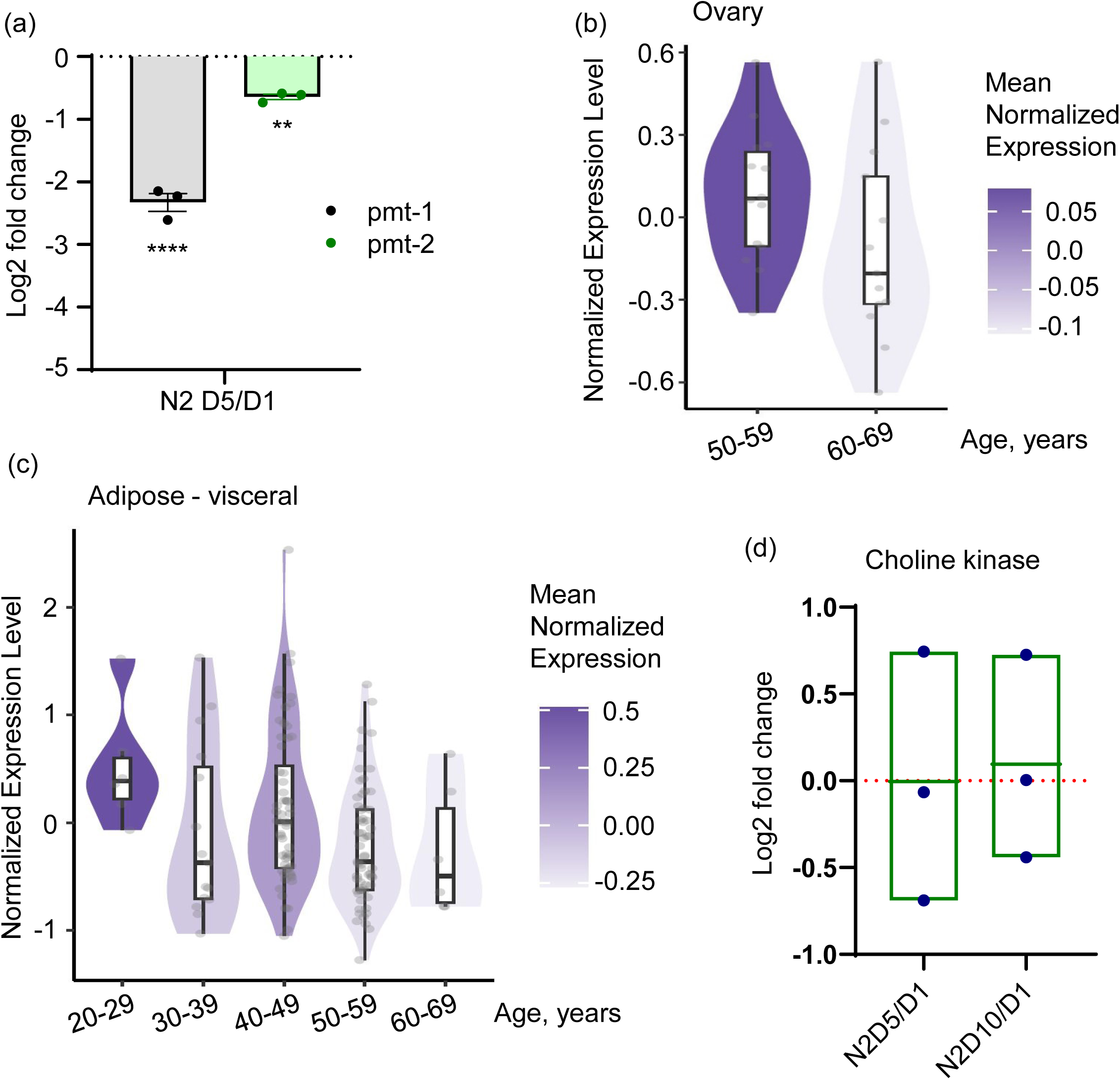
Expression of phosphatidylcholine producing enzymes in aging humans and *C. elegans*. **(a)** Proteomics samples were collected as described in Figures 1a-b and relative levels (expressed as Log 2 fold change) of PMT-1 and PMT-2 proteins in post-reproductive (AD5 vs AD1) WT animals are shown. All relevant calculations can be found in Table S4. **(b-c)** The expression of human PEMT gene in the ovary **(b)** and visceral adipose tissue **(c)** at different age was tested using the GTEx dataset (v8) as described in Figure 7b. Normalized expression levels represent individual log2 transformed quantile normalized Transcripts Per Million (TPM) values corrected for sex and death circumstances using a linear model; median normalized PEMT expression in each age group is color coded and complete data can be found in Tables S29 and S30. **(d)** Proteomics samples were collected as described in Figures 1a-b and relative expression (Log 2 fold change) of choline kinase subunits was measured in AD5 versus AD1 and AD10 versus AD1 WT animals. Each dot represent one protein and median values are shown. All relevant expression values can be found in Table S25.

**Figure S4.**
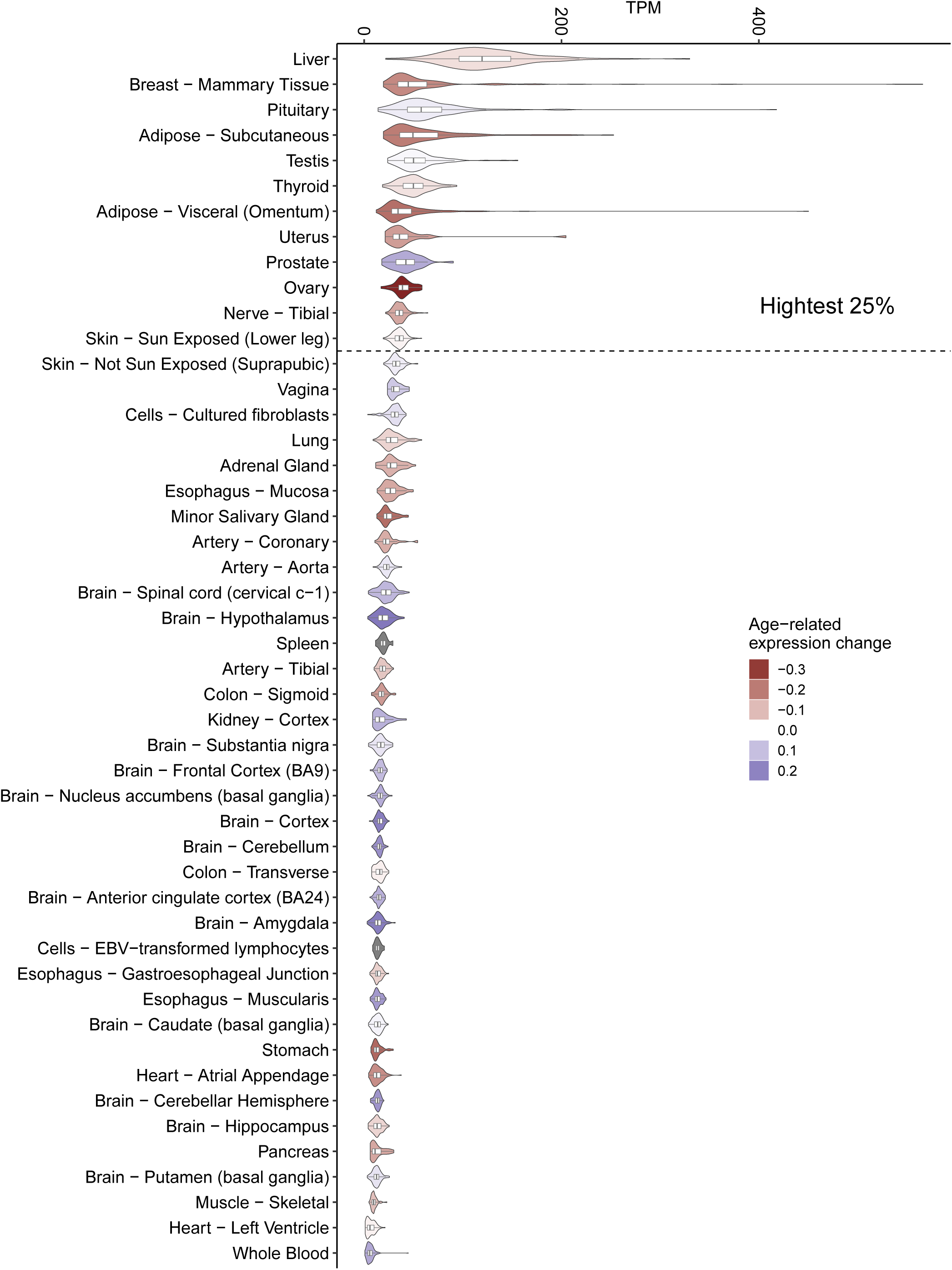
Expression of PEMT across human tissues and age groups. The expression of human PEMT gene across tissues at different age was tested using the GTEx dataset (v8) as described in Figure 7b. Relative PEMT expression in different organs is depicted as transcript per million (TPM), and highest expressing tissues are outlined as top 25%. Expression changes associated with aging are color coded. The analysis was performed as described in the methods section, including code. All relevant values can be found in Table S26.

**Figure S5.**
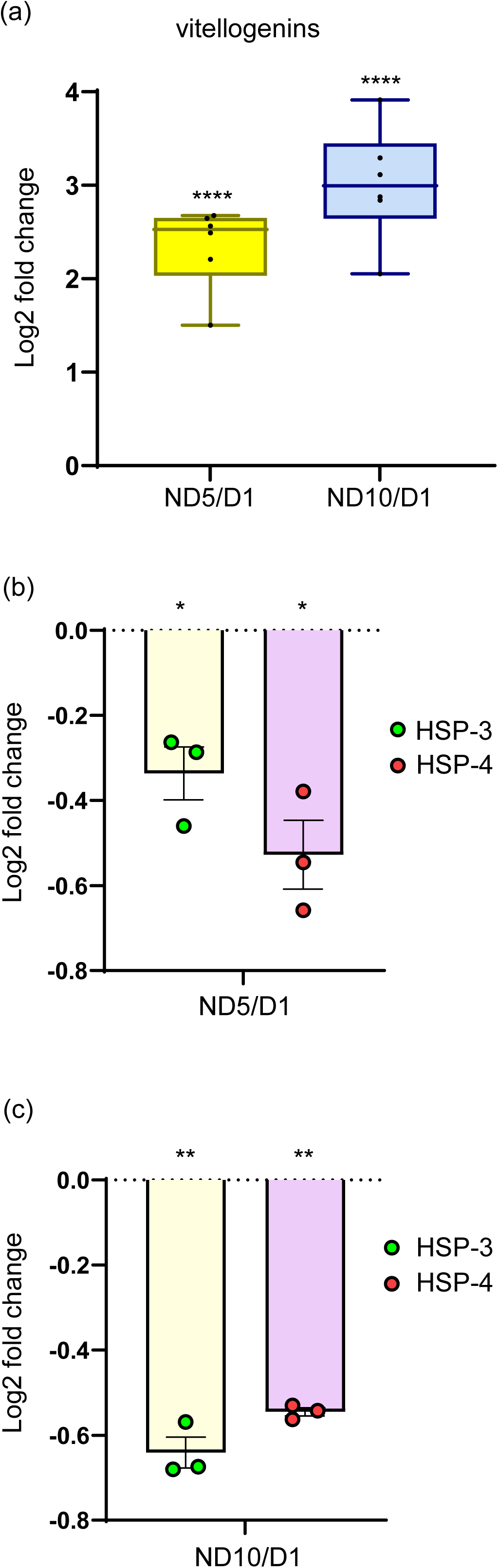
Elevated abundance of lipid droplets doesn’t coincide with the activation of UPR^ER^ in old nematodes. Proteomics samples were collected as described in Figures 1a-b. **(a)** Box plots depicting relative expression (Log 2 fold change) of vitellogenins in AD5 and AD10 versus AD1 WT nematodes are shown. Each dot represents one protein, median expression values are shown inside each box and whiskers highlight minimal and maximal values within each tested group. Statistics was assessed by one sample t-test and all relevant expression values can be found in Table S24. Relative expression (Log 2 fold change) of HSP-3 and HSP-4 ER chaperones is shown between AD5 and AD1 **(b)** and AD10 and AD1 **(c)** WT animals. Each dot represents one replica sample, mean and SEM values are presented and unpaired t-test was used to compute *p*-values. All relevant expression values and calculations can be found in Table S4. *-*p*<0.05; **-*p*<0.01; ****-*p*<0.0001, all values are two-tailed and exact *p* values can be found in the statistics Source data file.

**Figure S6.**
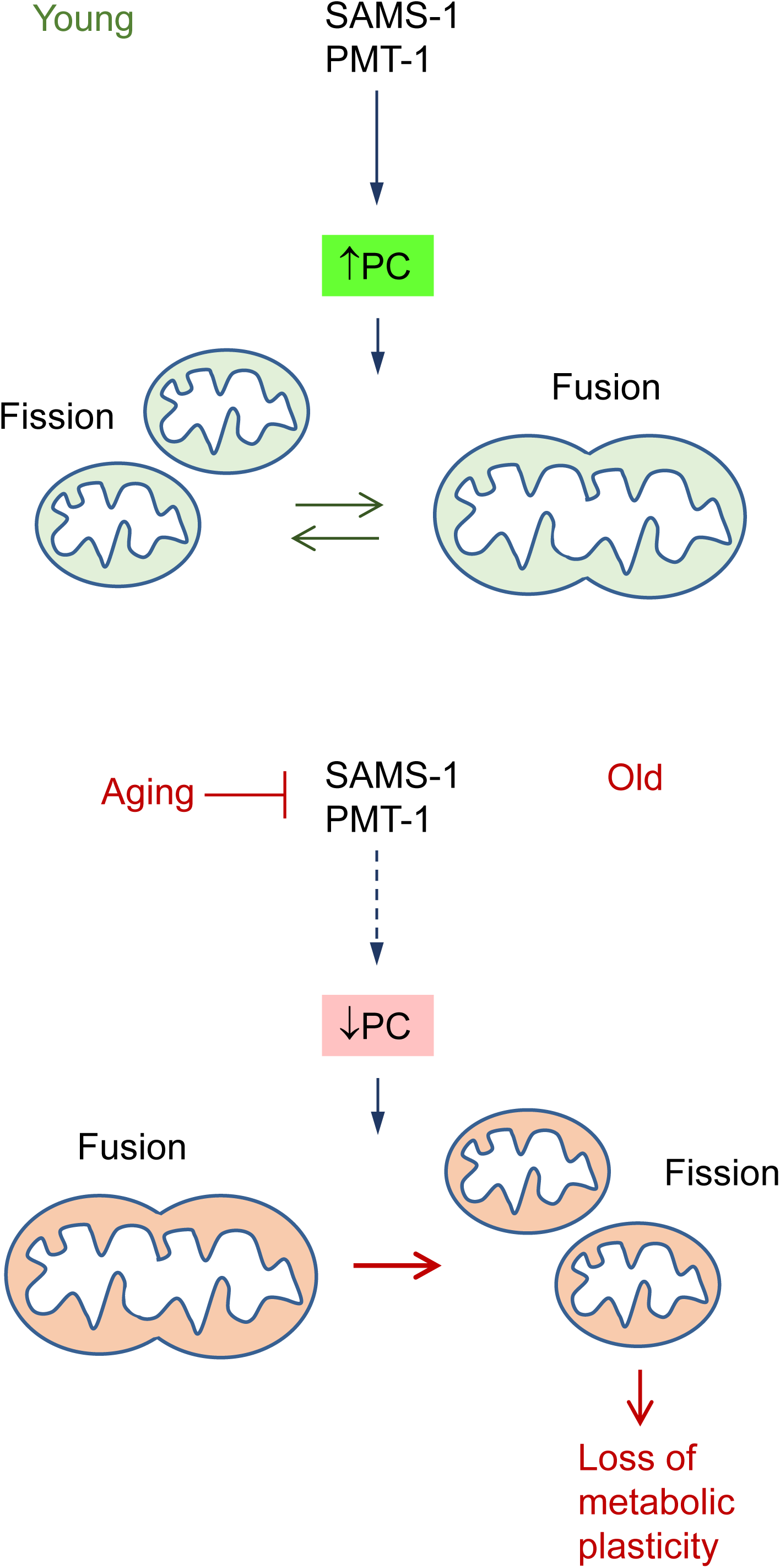
Aging-triggered decline of SAM-dependent phosphatidylcholine synthesis impairs mitochondrial fusion contributing to aging-associated mitochondrial dysfunction and loss of metabolic plasticity in late life. The schematic mechanistic model is presented; PC – phosphatidylcholine.

## Materials and methods

### C. elegans strains

The *C. elegans* strains were obstained from the Caenorhabditis Genetics Center (CGC) of the University of Minnesota, which is funded by NIH Office of Research Infrastructure Programs (P400D010440), www.cgc.umn.edu. Hermaphrodite nematodes were used in all tests, and maintained under standard conditions at 20°C unless otherwise noted. We used the following strains: N2 Bristol (wild-type), MQ130 *clk-1(qm30)* III, MQ887 *isp-1(qm150)* IV, SJ4103: *zcIs14[myo3p::gfp(mito)]*, SJ4100: *zcIs13[hsp-6::gfp]*.

### Bacterial strains

Worms were fed with OP50 *E. coli* obtained from CGC. Gene knockdowns were performed by feeding HT115 (DE3) L4440 *E.coli* obtained from ORF (*sams-1* RNAi) or Ahringer *C.elegans* RNAi collections (*mip-1* RNAi). *pmt-1* RNAi was kindly provided by Prof. O. Klotz (Friedrich Schiller University of Jena). Growth conditions for both OP50 and RNAi bacteria were as described in (Espada et al., 2020).

### Treatment with drugs and supplements

For worms, SAM (Hanoju) was extracted from capsules, dissolved in Milli Q® water, filtered with 0,22 µm filter to remove microcellulose impurities contained in the capsules, and applied on top of the bacterial lawn to reach a final concertation of 10 mM, considering the 10 ml NGM agar volume. Phosphatidylcholine (PC) concentration and application was chosen according to a previous report (Kim et al., 2019). Briefly, PC was dissolved in EMPROVE®exp ethanol (Merck) and the final concentration per NGM plate was chosen as 0.1% of ethanol and 100mg/L of PC (Sigma-Aldrich). Higher concentrations of ethanol promoted significant mitochondrial fragmentation in pilot tests. Both SAM and PC supplemented plates were kept under light-protective conditions. Choline (Sigma), was given to worms by addition to the NGM agar. In cell culture tests, succinate (Oroboros, MitoKit-CII) choline and metformin (M2009 TCI) were added directly to culture media.

### Lifespan assay

Lifespan assay was performed as described in (Espada et al., 2020).

### Imaging and quantification

Live fluorescence microscopy and image analyses were performed as in Espada et al., 2020 with some modifications. In testing effects of gene inactivation on mitochondrial morphology and stress levels, *zcIs14[myo3p::gfp(mito)]* and *zcIs13[hsp-6::gfp]* reporter strains were fed with respective RNAi or EV control bacteria from L1 stage and until imaging on adulthood day 2 (AD2). Drug and vehicle treatments were delivered from L4 stage and until imaging at AD2. In testing the effect of choline supplementation on mitochondrial aging, worms were grown under standard conditions from L1 stage and until adulthood day 1 (AD1) followed by choline or control exposure and imaging at indicated times. Mitochondrial morphology was assessed in the body wall muscles of the tail area in the *zcIs14[myo3p::gfp(mito)]* reporter strain. x63 magnification, oil immersion and temperature-controlled environment (15-17°C) were applied. The % of tubular, intermediate, fragmented and very fragmented mitochondria per worm was calculated. Mitochondrial stress was measured using *zcIs13[hsp-6::gfp]* reporter strain and x100 magnification. Each individual worm perimeter was encircled, and mean fluorescence was measured followed by background subtraction in Fiji - ImageJ software. Area of each worm calculated using Fiji-Image J was used to compare the size of the worms when necessary. In all cases, at least 20 worms per condition were used in 3 replicas (individual plates) and all experiments were repeated at least 3 times. For representative snapshots, equally scaled images were used.

### Quantitative PCR with reverse transcription

Total RNA was isolated from worms for cDNA synthesis, which was used for real time quantitative PCR, with data analysed using the ΔΔCt method (Livak & Schmittgen, 2001). Briefly, 60 worms at L4 stage were washed with M9 buffer, lysed in TRIzol ™ (Sigma-Aldrich), homogenized using ceramic beads (biostep GmbH) and frozen. Total RNA was isolated using 1-bromo-3-chloropropane (Merck) and the Analytik Jena Kit (#845-KS-2040050), followed by treatment with DNase I RNase free (Thermo Scientific™) and RiboLock RNAse Inhibitor (Thermo Scientific™). First-strand cDNA synthesis was performed from 0,5 µg RNA using SuperScript™ III Reverse Transcriptase kit (Invitrogen). Real time PCR was carried out using Bio-Rad SsoAdvanced™ Universal SYBR® Green Supermix 2x (Bio-Rad) and CFX 96™ Real-Time System with C1000 Touch™ Thermal Cycler. Mean value of 3 housekeeping genes - actin, tubulin and laminin, were used as a reference.

### Sample preparation for proteomics analysis

Worms (700 L1s grown and taken at indicated age) were washed 5 times with M9 Buffer (3 times in 10 mL, 2 times in 5 mL of buffer to remove eggs and progeny) and collected directly into bead-beating tubes to avoid losses during transfer. Lysis buffer (1x) was 1 % SDS, 100 mM HEPES, pH 8, 100 mM DTT, 1mM EDTA. Samples were bead-beaten in a Precellys bead-beater at 6000 rpm, 2×20 sec, 30 sec interval at 4 °C. Samples were centrifuged (1000 g, 3 min) and the supernatant pipetted into fresh 0.5 mL Eppendorf tubes®. Samples were then sonicated using a Bioruptor (60 sec ON/30 sec OFF, 10 cycles, high intensity at 20 °C) (Diagenode, Belgium), then heated to 95 °C for 10 min, before a repeat set of sonication cycles as before. Alkylation to block cysteines was carried out with iodoacetamide (15 mM final concentration, 30 min, dark, room temperature). Protein precipitation was carried out on an estimated 25 μg amount of each sample using 4x sample volume of ice-cold acetone, and samples were left at -20 °C overnight. The following day, samples were centrifuged (20800 g, 30 min, 4 °C), supernatant carefully removed, and protein pellets washed twice with ice cold 80 % acetone/20 % water (300 μL, 10 min centrifugation (as above). After removal of the second wash, pellets were air-dried before resuspension in the digestion buffer (3 M urea in 0.1 M HEPES, pH 8; LysC (1:100 enzyme: protein ratio), then incubated for 4 h at 37 °C. The samples were diluted 1:1 with Milli-Q® water (to reach 1.5 M urea) and incubated with trypsin (1:100 enzyme: protein) for 16 h at 37 °C. The digests were then acidified with 10 % trifluoroacetic acid and then desalted with Waters Oasis® HLB μElution Plate 30 μm (Waters Corporation, Milford, MA, USA) in the presence of a slow vacuum. In this process, the columns were conditioned with 3 x 100 μL solvent B (80 % acetonitrile; 0.05 % formic acid) and equilibrated with 3 x 100 μL solvent A (0.05 % formic acid in Milli-Q® water). The samples were loaded, washed 3 times with 100 μL solvent A, and then eluted into PCR tubes with 50 μL solvent B. The eluates were dried down with the speed vacuum centrifuge prior to resuspension in MS Buffer (5 % acetonitrile, 95 % Milli-Q® water, with 0.1 % formic acid) for the MS analyses.

### Proteomics data aquisition in data dependent (DDA) and independent (DIA) modes

Data were acquired on a QE-HFX MS (Thermo), connected to an M-Class nanoAcquity (Waters). The outlet of the analytical column was coupled directly to the mass spectrometer using the Proxeon nanospray source. The trapping column was nanoAcquity Symmetry C18, 5 μm, 180 μm x 20 mm and the analytical column was nanoAcquity BEH C18, 1.7 μm, 75 μm x 250 mm. Solvent A was water, 0.1 % formic acid, and solvent B was acetonitrile, 0.1 % formic acid. Approximately 1 μg of each of the samples (reconstituted at 1 μg/ μL and spiked with HRM kit peptides (Biognosys AG, Switzerland)) were injected for LC-MS with a constant flow of solvent A, at 5 μL/min, in trapping mode. The trapping time was 6 minutes. Peptides were eluted via the analytical column with a constant flow of 0.3 μL/min. During the elution step, the percentage of solvent B increased in a non-linear fashion from 0 % to 40 % in 120 minutes. Total runtime was 145 minutes, including clean-up and column re-equilibration. The peptides were introduced into the mass spectrometer via a Pico-Tip Emitter 360 μm OD x 20 μm ID; 10 μm tip (New Objective) and a spray voltage of 2.2 kV was applied. The capillary temperature was set at 300 °C. The ion funnel RF was set to 40 %. DDA data were acquired on a subset of samples from all conditions, with the following settings: Full scan MS spectra with mass range 350-1650 m/z were acquired in profile mode in the Orbitrap with resolution of 120000. MS1 fill time was 20 ms, AGC target 3 x 106. Top N was used (=15) and the intensity threshold was 4 x 104. Normalized Collision Energy (NCE) with HCD was set to 27 % and a 1.6 Da window was used for quadrupole isolation. MS2 data were acquired in profile mode from 200 – 2000 m/z. MS2 fill time was 25 ms or an AGC target of 2 x 105. Only 2-5+ charge states were selected for MS/MS. Dynamic exclusion was 20 s, and the peptide match ‘preferred’ option was selected. For the DIA data, LC conditions remained unchanged. Full scan MS spectra with mass range 350-1650 m/z were acquired in profile mode in the Orbitrap with resolution of 120000.The default charge state was set to 3+. The filling time was set at maximum of 60 ms with limitation of 3 x 106 ions. DIA scans were acquired with 34 mass window segments of differing widths across the MS1 mass range. HCD fragmentation (stepped normalized collision energy; 25.5, 27, 30 %) was applied and MS/MS spectra were acquired with a resolution of 30000 with a fixed first mass of 200 m/z after accumulation of 3 x 106 ions or after filling time of 40 ms (whichever occurred first). Data were acquired in profile mode. For data acquisition and processing of the raw data Xcalibur 4.0 (Thermo Scientific) was used in parallel with Tune version 2.9.

### Proteomics data analysis

For library creation, the DDA and DIA data were searched independently using the Pulsar search engine (Spectronaut™ (version 11.0.15038.17.27438); Biognosys AG, Zurich, Switzerland). Afterwards, the two libraries were merged to create a single DpD (<u>D</u>DA <u>p</u>lus <u>D</u>IA library). In both cases, data were searched against a species specific (*C. elegans*) UniProt database (as for the digest check) alongside the database of common contaminants. The data were searched with trypsin/P specificity and the following modifications: Carbamidomethyl (C) (Fixed) and Oxidation (M)/ Acetyl (Protein N-term) (Variable). A maximum of 2 missed cleavages was allowed. The identifications satisfied an FDR (false discovery rate) of 1 % on both peptide and protein level. All other settings were the defaults from Biognosys.

The resulting library contained 92001 precursors, corresponding to 5338 protein groups using Spectronaut™ protein inference. DIA data were then uploaded and searched against this spectral library in Spectronaut™. Relative quantification was performed in the software for each pairwise comparison using the replicates from each condition. Spectronaut™ ran a pairwise comparison at the peptide level and then summarised it at the protein level. Differences in protein abundances were statistically determined by Spectronaut™ software using the Student’s t-test and multiple testing correction algorithm described by Storey (2002). Contrast table (candidates table) and protein matrix table were then exported for further data visualization analysis using in-house scripts with R-studio (version 1.0.153). The further used for analysis here Q value represents the false discovery rate for multiple comparisons.

### Data availability

The mass spectrometry proteomics data have been deposited to the ProteomeXchange Consortium (Deutsch et al., 2020) via the PRIDE (Perez-Riverol et al., 2019) partner repository available at http://www.ebi.ac.uk/pride, with the dataset identifier PXD024180.

### Identification of relevant proteome changes

Relative quantification was performed in Spectronaut for each pairwise comparison among N2 and mito-mutants at AD1, AD5 and AD10 using the triplicate samples from each condition. The data (candidate table with log2 (fold changes) and Q values for each pairwise comparison, and raw intensities when applicable) were exported to Excel for further analyses presented as Venn diagrams, pie charts, bar charts and boxplots. Visualization was performed using Graph Prism 8.0.0.

The Venn diagrams compare lists of genes differentially expressed in N2 and mitomutants at AD1, AD5 and AD10. The online tool http://bioinformatics.psb.ugent.be/webtools/Venn/ was used to create the diagrams from pairwise comparisons subsetted in respective supplementary tables. For gene set enrichment analysis of overlapping and non-overlapping protein lists, the WormCat 2.0 online tool was used (www.wormcat.com).

### Cell culture experiments

Cell culture maintenance, the LDH (lactate dehydrogenase) cytotoxicity assay, and mitochondrial membrane potential JC-1 assay, were performed according to (Espada et al., 2020).

### Human data analysis

PEMT expression and age-related variations across tissues were analyzed using the GTEx dataset (v8) (https://www.nature.com/articles/ng.2653) and preprocessed data from Donertas et al 2021 (https://www.nature.com/articles/s43587-021-00051-5). These data incorporated individuals with Hardy Scale scores 1 and 2, filtered out genes with a median TPM below 1, applied log2 transformation, and quantile normalized the dataset after correcting for sex and death circumstances using a linear model. Samples deviating more than 3 standard deviations from the first four principal components were considered outliers and excluded. Preprocessing scripts are available in Github https://github.com/mdonertas/aging_in_GTEx_v8. Age-related expression changes were calculated using Spearman’s correlation for individuals for tissue-age pairs with more than five samples. Tissues with high PEMT expression were determined by computing mean TPM values, following the method employed in GTEx data portal but restricting data to same individuals used in our main analysis (i.e. excluding slow death and ventilator cases and outliers). All resources, including code, processed data, and results, can be accessed at https://github.com/mdonertas/pc_synthesis_human_aging.

Code for PEMT analysis is at https://github.com/mdonertas/pc_synthesis_human_aging. Code for GTEx data preprocessing is at https://github.com/mdonertas/aging_in_GTEx_v8.

### Statistical analysis

The results are expressed as mean + SEM in most cases unless stated otherwise. Statistical analyses were performed in R, GraphPad Prism 8.0.0 (GraphPad Software Inc.) and Excel. Respective statistical tests are described in each figure legend and exact *p* values and *n* numbers are provided in the Statistics summary file for each experiment.

